# Hypovolemia Evokes Conserved Inverse Neurovascular Coupling in the Supraoptic Nucleus Independent of Heart Failure

**DOI:** 10.1101/2025.11.22.689917

**Authors:** Ranjan K. Roy, Jessica A. Filosa, Javier E. Stern

**Affiliations:** Neuroscience Institute, Georgia State University; Center for Neuroinflammation and Cardiometabolic Diseases, Georgia State University; Department of Physiology, Augusta University, Augusta GA USA

## Abstract

Vasopressin (AVP) neurons in the hypothalamic supraoptic nucleus (SON) are activated by systemic challenges that threaten fluid balance. Beyond their classical activity-dependent release of their neuropeptide cargo into the systemic circulation, these neurons also release AVP somatodendritically, enabling local modulation of neuronal excitability and vascular tone. We previously showed that a systemic salt challenge triggers inverse neurovascular coupling (iNVC) in the SON, in which activity-dependent dendritic AVP release induces parenchymal arteriole vasoconstriction and local hypoxia. In rats with heart failure (HF), however, the polarity of this salt-evoked response is reversed: microglia-derived adenosine acting on A2A receptors overrides an enhanced AVP-mediated vasoconstriction, producing net vasodilation. Still, whether AVP activation by non-osmotic stimuli engages similar neurovascular mechanisms is unknown. Here, we examined whether hypovolemia induced by intraperitoneal polyethylene glycol (PEG) evokes comparable vascular responses in control and HF rats. PEG produced a sustained rise in plasma protein concentration and vasoconstriction of SON parenchymal arterioles in both control and sham rats. In HF rats, PEG still induced vasoconstriction at 60 min, but vascular diameters returned to baseline by 90 min despite persistent hypovolemia. These findings indicate that hypovolemia engages a conserved AVP-mediated iNVC program that remains largely intact in HF, and that the previously described polarity reversal during HF is stimulus-specific, emerging during osmotic but not hypovolemic activation of AVP neurons.

## INTRODUCTION

Arginine vasopressin (AVP) neurons in the hypothalamic supraoptic (SON) and paraventricular (PVN) nuclei play a central role in regulating fluid and electrolyte homeostasis. In response to systemic challenges such as increased plasma osmolality or decreased circulating volume, AVP neurons increase their electrical activity and release AVP into the circulation to promote water conservation and vascular regulation(1). In addition to their classical systemic output, accumulating evidence demonstrates that these neurons also release AVP somatodendritically within the hypothalamus, where locally acting AVP can shape neuronal excitability, modulate synaptic integration, and influence broader hypothalamic network function(2). A growing body of work indicates that dendritically released AVP can also regulate local vascular dynamics. In this sense, we recently showed that a systemic salt challenge, a potent activator of AVP neurons, elicits a form of inverse neurovascular coupling (iNVC) in the SON(3). This unconventional form of NVC is mediated by the activity-dependent somatodendritic release of AVP, which induces vasoconstriction of SON parenchymal arterioles, generating a progressive local hypoxic milieu that further enhances AVP neuronal excitability(3). This positive feedback mechanism reveals a previously unrecognized coupling between neuropeptide release and microvascular tone in deep hypothalamic circuits.

Unexpectedly, we also found that the polarity of this salt-evoked NVC response is altered in rats with heart failure (HF)(4), a condition in which the AVP system is overly activated (5-9), contributing in turn to morbidity and mortality in this disease (10-12). We found that while AVP can still evoke vasoconstriction in HF rats (which is, in fact, of greater magnitude than in control conditions), this effect is overridden by a dominant vasodilatory signal mediated by adenosine acting on A2A receptors(4). Pharmacological and mechanistic evidence suggested that this adenosine signal likely originates from microglia, revealing a stimulus- and disease-specific shift in the balance of vasoactive modulators within the SON during HF.

All these previous observations were made using a hyperosmotic stimulus to activate AVP neurons. Still, whether AVP neurons engaged by a distinct homeostatic challenge, such as hypovolemia, generate similar neurovascular responses remains unknown. Hypovolemia activates AVP neurons via mechanistically distinct afferent pathways compared with osmotic stimulation (13), raising the question of whether the resulting NVC pattern is conserved, or conversely, whether it is stimulus-specific. Addressing this gap is essential for determining whether iNVC represents a general property of AVP neuronal activation or a response restricted to the osmotic axis.

In this study, we examined whether hypovolemia elicits comparable neurovascular dynamics in the SON of healthy and HF rats. We used intraperitoneal polyethylene glycol (PEG), a well-established and robust approach to induce hypovolemia and activate AVP neurons(14, 15). Using in vivo two-photon imaging of SON parenchymal arterioles in control, sham, and HF rats, we assessed the temporal dynamics and polarity of vascular responses following PEG treatment.

Our findings reveal that PEG-induced hypovolemia evokes vasoconstriction of SON parenchymal arterioles in control rats, consistent with the iNVC previously observed with salt. However, unlike the salt response, the polarity of the vascular response remains unchanged in HF, indicating that the reversal of NVC polarity is not a universal consequence of AVP neuronal activation, but rather a stimulus-specific feature of the salt stimulation. These results identify hypovolemia as a condition in which AVP-mediated iNVC is preserved despite the HF pathology, underscoring the stimulus-specific tuning of hypothalamic neurovascular mechanisms.

## MATERIALS AND METHODS

### Animals

Male and female heterozygous transgenic eGFP-AVP Wistar rats (250-400gm) were used for all experiments (16). Rats were housed in cages (2 per cage) under constant temperature (22 ± 2°C) and humidity (55 ± 5%) on a 12-h light cycle (lights on: 08:00-20:00). All performed experiments were approved by the Georgia State University Institutional Animal Care and Use Committee (IACUC) and carried out in agreement with the IACUC guidelines. At all times, animals had *ad libitum* access to food and water, and every effort was made to minimize suffering and the number of animals used in this study.

### Heart failure surgery and echocardiography

To induce heart failure in rats, a coronary artery ligation surgery was performed as previously described (17). Anesthesia was induced using 5% isoflurane mixed with O_2_ (100% O_2_, 1L/min). Optimum anesthesia was assessed by the absence of limb withdrawal reflex to a painful stimulus (hind paw pinch). Rats were then intubated for mechanical ventilation. During the surgery, anesthesia was adequately maintained using 2-3% isoflurane delivered by a vaporizer machine mixed with O_2_ (100% O_2_, 1L/min). With aseptic precaution, a small incision was made in the third intercostal space, and a retractor was applied to retract the 3rd and 4^th^ ribs. The pericardium was ruptured, and the heart was exteriorized. A ‘Prolene 6-0’ suture (Ethicone, USA) was inserted just beneath the left atrial base along the interventricular septum to make a loop around the main diagonal branch of the left anterior descending (LAD) coronary artery. Finally, myocardial infarction was induced by ligating the LAD coronary artery. Occlusion of the LAD coronary artery results in a pale discoloration of the left ventricle, which serves as a visual confirmation of the ischemic left ventricle following LAD coronary artery ligation. Sham animals underwent a similar aseptic surgical procedure except for left coronary artery ligation. Four to five weeks after the surgery, we performed transthoracic echocardiography (Vevo 3100 systems; Visual Sonics, Toronto, ON; Canada) under light isoflurane (2-3%) anesthesia to assess the ejection fraction (EF) and confirm the development of HF. We obtained the left ventricle’s internal diameter and the diameter of the ventricle’s posterior and anterior walls in the short-axis motion imaging mode to calculate the EF. Myocardial infarction surgery typically results in a wide range of functional HF, as determined by EF measurements. Rats with EF<40% were considered as HF (Sham 84.34 ± 3.09 %, HF 30.44 ± 6.19%; n= 6 and 10 in sham and HF rats, respectively, see Table 1 for additional echocardiographic parameters). Rats that underwent HF surgery but did not develop HF or displayed an EF>40% were not included in the study.

**Table 1.** Summary of assessed cardiac parameters in sham and heart failure (HF) rats via echocardiography.

|  | **Sham**<br>**(n=6)** | **HF**<br>**(n= 10)** | **P value**<br>**(unpaired t-test)** |
| --- | --- | --- | --- |
| **Heart rate (BPM)** | 288.46 ± 25.67 | 292.27 ± 23.32 | P = 0.237 |
| **Stroke Volume (uL)** | 230.39 ± 27.22 | 140.72 ± 10.33 | P < 0.001 |
| **Ejection Fraction (%)** | 84.34 ± 3.09 | 30.44 ± 6.19 | P < 0.001 |
| **Fractional Shortening (%)** | 45.06 ± 4.12 | 17.33 ± 2.11 | P < 0.001 |
| **Cardiac Output (mL/min)** | 67.37 ± 5.53 | 41.07 ± 2.62 | P < 0.001 |
| **LV Mass (mg)** | 1813.67 ± 64.24 | 989 ± 46.09 | P < 0.001 |
| **Lvaw,d (mm)** | 3.67 ± 0.65 | 1.69 ± 0.47 | P < 0.001 |
| **Lvaw,s (mm)** | 3.88 ± 0.18 | 1.55 ± 0.18 | P < 0.001 |
| **Lvid,d (mm)** | 5.47 ± 0.56 | 11.67 ± 0.36 | P < 0.001 |
| **Lvid,s (mm)** | 4.35 ± 0.5 | 8.03 ± 0.11 | P < 0.001 |
| **Lvpw,d (mm)** | 4.23 ± 0.38 | 1.3 ± 0.73 | P < 0.001 |
| **Lvpw,s (mm)** | 3.55 ± 0.47 | 1.86 ± 0.27 | P < 0.001 |

### Surgery for *in vivo* 2-photon imaging of the rat supraoptic nucleus (SON)

A modified transpharyngeal surgical approach was used to expose the ventral surface of the brain containing the hypothalamic SON (3, 18). Briefly, rats were anesthetized by intraperitoneal injection of urethane (1.5 gm/Kg) (U2500, Sigma Aldrich, MO, USA). Upon cessation of the hindlimb withdrawal reflex, the trachea and left femoral vein were catheterized, and the rats were placed supine on a heating pad in a stereotaxic frame (Stoelting 51600 Lab Standard Stereotaxic Instrument). After the oral cavity was opened by splitting the mandibles, cautery (Thermal Cautery Unit®, Geiger Medical Technologies, IA, USA) and drilling were used to remove the molars, ventromedial aspect of the right temporal bone, the basisphenoid and presphenoid bones to expose the ventral surface of the hypothalamus and ventral aspect of the right temporal lobe. The meninges overlying the right SON were removed, and a stainless-steel ring (i.d. 3.6 mm) was placed on the surface of the brain with the junction of the internal carotid artery (ICA) and middle cerebral artery (MCA) at the center. All the drugs used were topically delivered through an MD probe (infusion rate 0.6ml/hour), which only penetrates a short distance inside the brain surface (19).

### In-vivo 2-photon imaging of the SON and surrounding vasculature

The microvasculature of the SON and the surrounding area was labelled by i.v. injection of Rhodamine 70 kDa (20 mg/ml, 200 nl/rat) (R9379, Sigma Aldrich, MO, USA). The exposed ventral surface of the brain containing the SON and its surrounding vasculature was imaged under a 2-photon microscope (Bruker, Billerica, MA) excited with a Ti: Sapphire laser (Chameleon Ultra II [Coherent, Santa Clara, CA]) tuned at 840 or 940 nm and scanned with resonant galvanometers. As previously characterized (3), the visually accessible area of the SON is typically located medially to the bifurcation of the ICA and MCA. For measurements of parenchymal artery (PA) diameter, we used a 4X (numerical aperture 0.13) objective (Olympus, Center Valley, PA, USA).

### Quantification of vessel diameter

Polyethylene Glycol (PEG) (5 g/Kg) was injected intraperitoneally (IP) in HF and control (sham) rats. To quantify changes in vessel diameter at baseline, 60, and 90 min post PEG injection, Z-stack acquisition protocol was used (20 images at 0.33 Hz, 30 µm interval up/down the Z-axis in the SON; 512 x 512 pixels). Analyses were performed with *ImageJ*, as previously described (3). Briefly, a Z-projected image was obtained, and vessel cross-section lines were manually drawn perpendicular to the two sides of the vessel wall. A profile of image brightness along the vessel’s cross-section was then obtained. The high contrast between the vessel wall and the brain parenchyma allows for the ready detection of the vessel wall edges, which appear as a sudden rise in basal fluorescence. The distance between the two fast-rising phases, measured just above the base of the fluorescence profile, was used to calculate the diameter in each frame of the z-stack.

### Measurement of plasma protein

Blood was collected from the femoral vein after the IP injection of PEG (5 g/Kg) (22, 23). Blood samples from the femoral vein were collected at 0, 15, 30, 45, 60, 75, and 90 min post-PEG injection and plasma was separated by centrifugation at 2000×g for 10 minutes in a refrigerated centrifuge. Protein concentration was measured from the plasma sample using a Nanodrop Ultra spectrophotometer (Thermo Fisher Scientific).

### Statistical analysis

All statistical analyses were performed using GraphPad Prism 10 (GraphPad Software, California, USA). The significance of differences was determined using paired or unpaired t-tests (two-sided in all cases), one-way or two-way repeated-measures (RM), or standard analysis of variance (ANOVA) with the post hoc Bonferroni’s multiple comparisons test. Results are expressed as means ± standard deviation (SD). Results were considered to be statistically significant if p < 0.05 and are presented as ^∗^ for p < 0.05, ^∗∗^ for p < 0.01, and ^∗∗∗^ for p < 0.0001 in the respective figures.

## RESULTS4

### Hypovolemia triggers vasoconstriction of SON parenchymal arterioles

To investigate whether hypovolemia-induced AVP neuronal activation(22) resulted in dendritic release of AVP within the SON and the concomitant inverse neurovascular coupling (iNVC) response, as we previously showed to be the case in response to a salt challenge (3), anesthetized rats with the SON exposed following a transpharyngeal surgery were positioned under a two-photon microscopy system. To induce hypovolemia, rats were intraperitoneally injected with polyethylene glycol (PEG) (5 g/Kg, I.P.) (22, 23). To confirm hypovolemia, we collected blood samples from the femoral vein at 0, 15, 30, 45, 60, 75, and 90 min post-PEG injection and subsequently measured plasma protein concentration using a Nanodrop spectrophotometer. As shown in ***Fig.1***, plasma protein concentration became significantly elevated starting at 30 min post-PEG, remaining elevated until 90 min post-PEG (one-way RM ANOVA; F= 16.60, p < 0.001; n = 4).

**Figure 1.**
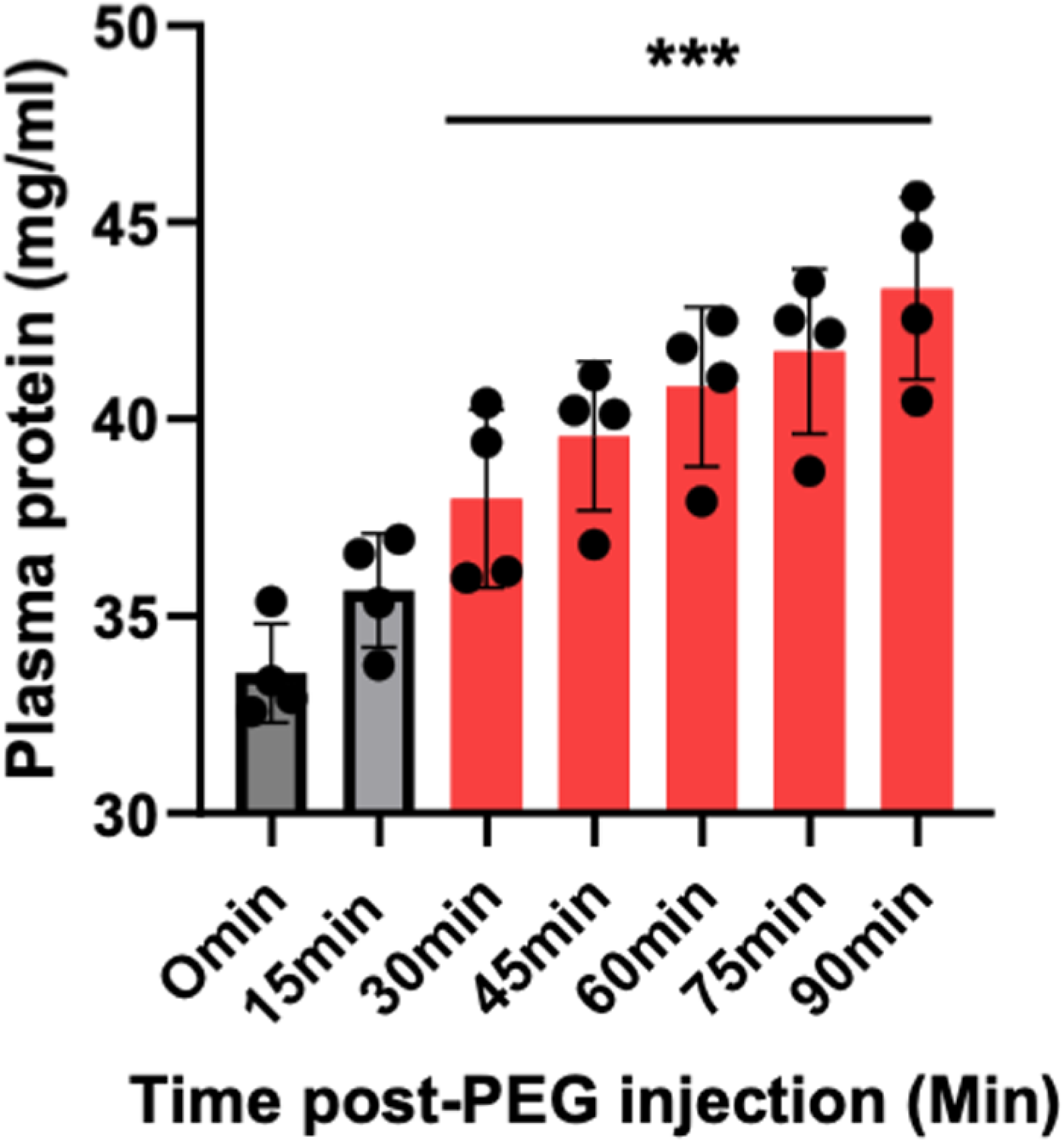
Plasma protein concentration increased after I.P. injection of PEG. Plasma protein concentration becomes significantly elevated starting at 30 min post-PEG, remaining elevated until the 90 min post-PEG tested (one-way RM ANOVA; F= 16.60, p < 0.001; n = 4).

Based on these results, we assessed vascular responses induced by hypovolemia at baseline and 60 minutes post-PEG. Similar to the salt-induced PA vasoconstriction (3), we found that PEG-induced hypovolemia resulted in a significant vasoconstriction at 60 min post-PEG (86.00 ± 2.86 μm at baseline, 76.97 ± 2.84 μm at 60 min post PEG injection, n=6, p<0.001, Bonferroni’s post hoc test (***Fig 2A,C***). Conversely, control saline (0.9% NaCl) I.P. injections failed to induce any change in SON PA diameters (84.11 ± 1.97 μm at baseline, 82.35 ±1.36 μm at 60 min post 0.9% NaCl injection, n=3, p=0.33 vs 60 min, p=0.67, Bonferroni’s post hoc test (***Fig 2B,C)***.

**Figure 2.**
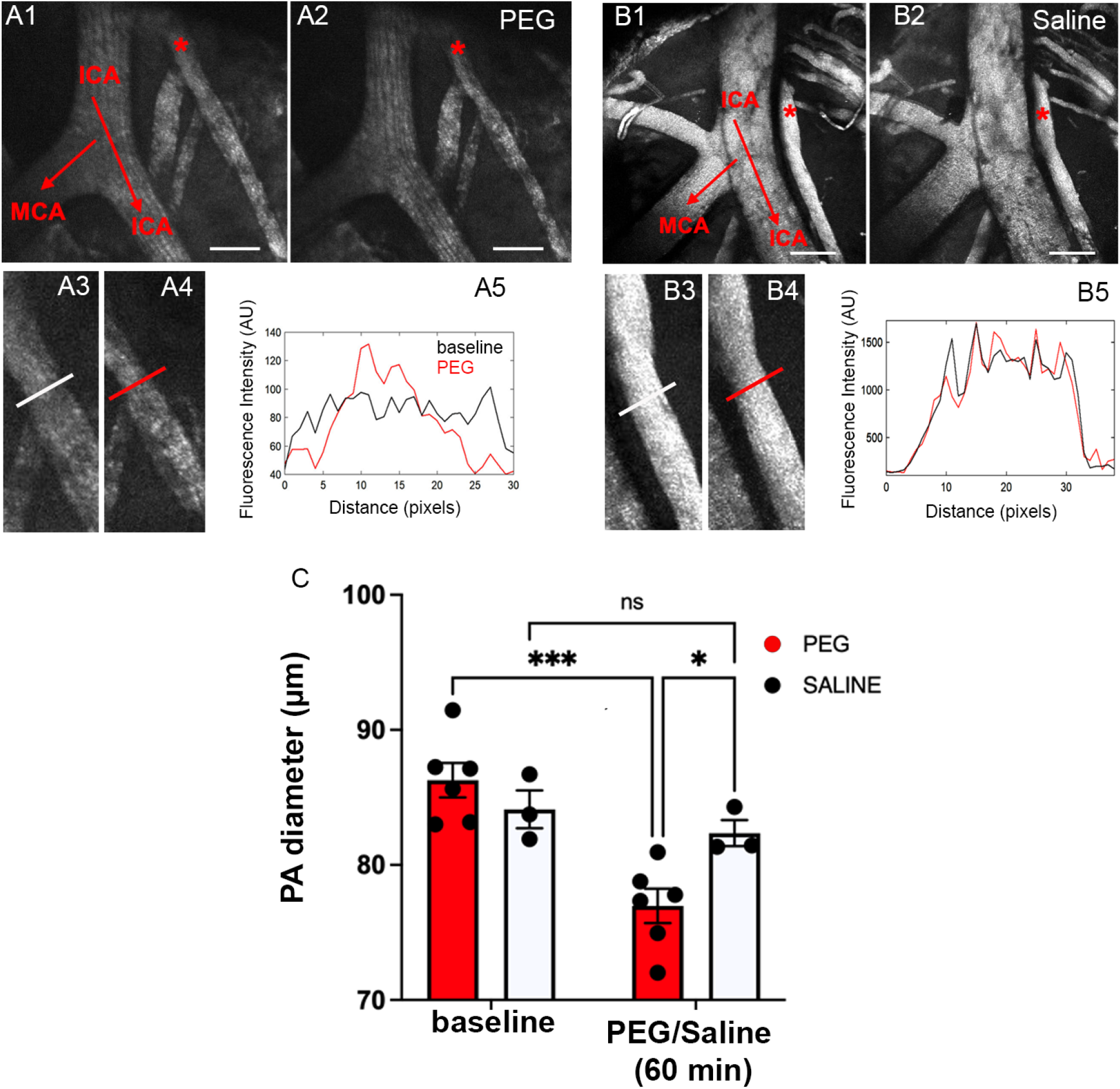
PEG evokes vasoconstriction of SON parenchymal arterioles (PA) in control rats. **(A)** Low magnification *in vivo* two-photon images of the SON following rhodamine 70 kDa administration (IV) before (A1) and 60 min after (A2) IP injection of PEG (5 g/Kg). A3 and A4 represent a magnified image of segments of a PA (demarcated as a red asterisk) in A1 and A2, respectively. A5, profiles of image brightness along the cross-section line of the PA vessel before (black) and 60 minutes post-PEG (red). Note the vasoconstriction evoked by PEG. **(B)** Low magnification *in vivo* two-photon images of the SON following rhodamine 70 kDa administration (IV) before (B1) and 60 min after (B2) IP injection of an equal volume of 0.9% NaCl). B3 and B4 represent a magnified image of segments of a PA (demarcated as a red asterisk) in B1 and B2, respectively. B5, profiles of image brightness along the cross-section line of the PA vessel before (black) and 60 minutes post-saline (red), showing lack of changes in PA diameter. **(C)** Bar graph showing the mean changes in PA diameter before and 60 min after IP PEG or saline injections. n=6 and 3 rats post-PEG and saline, respectively. *p< 0.05 and ***p<0.0001, Bonferroni’s post hoc test. Scale bars = 200μm

### The polarity of the hypovolemia-induced vascular response does not change in HF rats

Recently, we reported a reversal of the salt-induced iNVC in rats with HF, demonstrating salt-induced vasodilation, rather than vasoconstriction in this condition (24). To determine whether this was the case for hypovolemia, we repeated the PEG injections in sham and HF rats. In this case, we extended the recordings to 90 min post-PEG to determine whether effects persisted or changed over time. Similar to what we observed in control rats, PEG-induced hypovolemia resulted in PA vasoconstriction in both sham and HF rats (2-way ANOVA, F= 25.7, p < 0.0001). Thus, in sham rats, PEG resulted in vasoconstriction at both 60 and 90 min compared to baseline (p< 0.001 and p< 0.05, respectively, Bonferroni’s post-hoc test, n= 6 rats) ***(Fig. 3A,C)***. In HF rats, however, the vasoconstriction was observed only at 60 min post-PEG (p< 0.05, Bonferroni’s post-hoc test, n= 6 rats), returning to baseline levels at 90 min post-PEG (p= 0.16, Bonferroni’s post-hoc test, n= 6 rats) (***Fig.3B,C***). Taken together, these results indicate that hypovolemia-induced activation of AVP neurons leads to a similar iNVC as we observed in response to salt. Notably, the reversed polarity of this NVC response following activation of AVP neurons is observed only in response to salt, not to hypovolemia.

**Figure 3.**
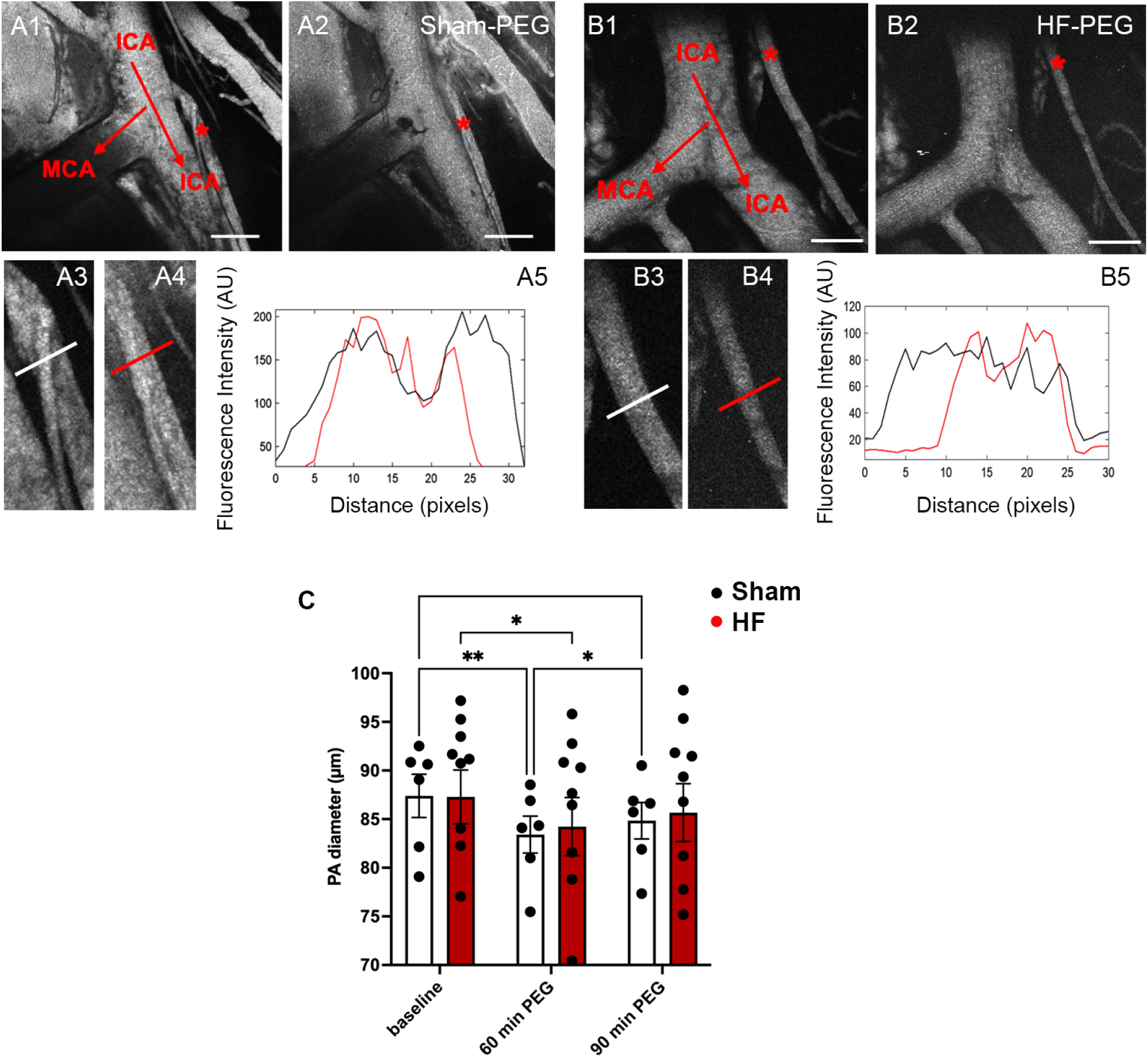
PEG-evoked vasoconstriction in SON parenchymal arterioles (PA) persists up to 90 min in sham rats but returns to baseline at 90min in HF rats. **(A)** Low magnification *in vivo* two-photon images of the SON in a sham rat following rhodamine 70 kDa administration (IV) before (A1) and 60 min after (A2) IP injection of PEG (5 g/Kg). A3 and A4 represent a magnified image of segments of a PA (demarcated as a red asterisk) in A1 and A2, respectively. A5, profiles of image brightness along the cross-section line of the PA vessel before (black) and 60 minutes post-PEG (red). Note the vasoconstriction evoked by PEG. **(B)** Low magnification *in vivo* two-photon images of the SON of an HF rat following rhodamine 70 kDa administration (IV) before (B1) and 60 min after (B2) IP injection of PEG (5 g/Kg). B3 and B4 represent a magnified image of segments of a PA (demarcated as a red asterisk) in B1 and B2, respectively. B5, profiles of image brightness along the cross-section line of the PA vessel before (black) and 60 minutes post-PEG (red). Note the vasoconstriction evoked by PEG. **(C)** Bar graph showing the mean changes in PA diameter before, 60 and 90 min pst-PEG in sham and HF rats (n=6 rats in each group). *p< 0.05 and **p<0.001, Bonferroni’s post hoc test. Scale bars = 200μm.

## DISCUSSION

In this study, we aimed to determine whether hypovolemia, a physiologically relevant non-osmotic stimulus of vasopressin (AVP) neurons, elicits neurovascular responses in the supraoptic nucleus (SON) comparable to those previously described during osmotic activation (3, 4). Specifically, we asked two main questions: (A) whether other systemic challenges that activate AVP neurons (e.g., hypovolemia) are sufficient to trigger inverse neurovascular coupling (iNVC) in control rats, and (B) whether the polarity of this response is similarly altered in heart failure (HF), as we previously demonstrated for salt-evoked osmotic stimulation (4).

Our main findings can be summarized as follows: (1) intraperitoneal polyethylene glycol (PEG), which induces a robust and sustained hypovolemic state, evoked a significant vasoconstriction of SON parenchymal arterioles in control rats, consistent with the iNVC previously observed during osmotic stimulation; (2) PEG-induced hypovolemia produced a comparable vasoconstrictive response in sham-operated animals, indicating that the iNVC response is preserved across both naive and surgically manipulated control groups; (3) In contrast to the polarity reversal we observed during salt stimulation in HF rats (4), hypovolemia did not reverse the neurovascular response in HF. Instead, HF animals displayed an initial vasoconstriction at 60 min post-PEG, with vascular diameter returning to baseline by 90 min, despite persistent hypovolemia.

Together, these findings demonstrate that hypovolemia engages a conserved AVP-mediated iNVC program in the SON, and that the polarity reversal we previously identified in HF is stimulus-specific, emerging during osmotic but not hypovolemic activation of AVP neurons.

### Hypovolemia, similar to an osmotic challenge, induced an inverse neurovascular coupling response in the SON

We previously showed that osmotic activation of AVP neurons via a systemic salt challenge leads to robust somatodendritic release of AVP within the SON, which in turn induces vasoconstriction of parenchymal arterioles and a progressive decline in local pO_2_, a form of inverse neurovascular coupling (iNVC)(3). This response aligns with previous evidence that osmotic stimulation increases AVP neuron firing and drives both systemic secretion and somatodendritic release of the peptide (1, 2).

However, whether physiologically distinct AVP-activating stimuli can similarly evoke iNVC has remained unknown. Although both osmotic stress and hypovolemia activate AVP neurons, the underlying mechanisms differ substantially. Osmotic stimuli can directly depolarize AVP neurons through intrinsic osmosensitive, TRPV1-like channels (25). In contrast, hypovolemia engages a separate set of afferent pathways, mainly arising from baroreceptor and volume receptor inputs relayed through the brainstem (26). These distinctions raise important questions about whether hypovolemia should trigger local dendritic release of AVP and, consequently, iNVC, to the same extent as osmotic activation. Indeed, to our knowledge, no prior studies have demonstrated that hypovolemia can induce dendritic release of AVP.

Although we did not directly measure dendritic release in this study, our findings show that hypovolemia reliably evokes iNVC in the SON of control rats. Based on our previous work, which established AVP as the primary mediator of salt-evoked iNVC, it is likely that a similar AVP-dependent mechanism underlies the hypovolemia-induced vasoconstriction reported here. Direct demonstration of hypovolemia-evoked dendritic AVP release will be an important future step. Collectively, our results suggest that hypovolemia is capable of driving local, AVP-dependent vascular responses in the SON, and together with osmotic stimulation, indicate that increased AVP neuronal activity per se may be sufficient to engage iNVC, irrespective of the specific physiological pathway through which AVP neurons are activated.

### Absence of polarity reversal in the hypovolemia-evoked SON neurovascular response during heart failure

We recently showed that during osmotic stimulation, the polarity of the neurovascular response in the SON is fundamentally altered in rats with heart failure (HF) (4). Salt-evoked activation of AVP neurons in HF rats no longer produced the expected vasoconstriction but instead triggered a dominant vasodilatory response mediated by microglia-derived adenosine acting on A2A receptors. Importantly, this vasodilation did not reflect a loss of AVP signaling; rather, it masked an underlying AVP-mediated vasoconstriction that became unmasked only when A2A or P2Y12 receptors were blocked (4). Thus, osmotic stimulation in HF recruits a second, overriding vasoactive mechanism that reversed the polarity of the net vascular response.

In contrast, in the present study, we found that hypovolemia-evoked activation of AVP neurons did not recruit this alternative vasodilatory pathway in HF. PEG-induced hypovolemia produced vasoconstriction at 60 min, consistent with a preserved iNVC driven by AVP, but unlike the osmotic stimulation, no vasodilatory reversal was observed, and vascular tone returned to baseline levels by 90 min. This difference raises two possible mechanistic considerations. First, the presence of a salt-specific vasodilatory mechanism in HF suggests that osmotic stimulation engages cellular elements within the SON, most notably microglia, that hypovolemia does not. High salt has been shown to directly activate microglia and promote pro-inflammatory signaling in the hypothalamus (27, 28) and other brain regions as well (29, 30). Such priming (which would be absent in the hypovolemic challenge) could facilitate rapid extracellular adenosine production via microglial ATP release and ectonucleotidase activity, leading to A2A receptor-mediated vasodilation. This differential microglial engagement offers a plausible explanation for why osmotic (but not hypovolemic) activation of AVP neurons recruits a vasodilatory override mechanism in HF.

Second, the overall hemodynamic and fluid/electrolyte state of HF itself may differentially constrain or modify the response to hypovolemia. HF is characterized by chronic fluid retention, elevated venous pressure, interstitial edema, and altered effective circulating volume(31, 32). This state produces a persistent activation of neurohumoral pathways, which may partially desensitize or saturate components of the hypovolemic reflex arc. As a result, an acute hypovolemic stimulus in this condition may be less effective in driving a sustained central response in HF. This could explain why PEG-induced vasoconstriction in HF rats was transient: iNVC was initially present, indicating that AVP neurons were activated and capable of inducing vasoconstriction, but the response in HF (but not sham rats) dissipated by 90 min despite persistent elevations in plasma protein concentration. Such attenuation could reflect impaired volume sensing, altered baroreceptor gain, or ceiling effects imposed by the chronically dysregulated volume state characteristic of HF.

Taken together, these observations suggest that hypovolemia, unlike osmotic stimulation, does not engage the adenosine signaling in the SON of HF rats, allowing the underlying AVP-mediated vasoconstriction to predominate. At the same time, the blunted temporal profile of the hypovolemic response in HF suggests that chronic alterations in fluid status may limit the ability of additional hypovolemic signals to sustain neuronal and vascular responses. Thus, the polarity and temporal dynamics of NVC in HF appear to be highly stimulus-dependent, shaped both by the intrinsic properties of the activating signal and by the altered physiological milieu imposed by HF.

### Study limitations and future directions

Several limitations of the present study should be acknowledged. First, although our findings strongly suggest that PEG-induced hypovolemia evokes inverse neurovascular coupling (iNVC) via local somatodendritic AVP release, we did not directly measure dendritic AVP release during hypovolemic stimulation. Given that no prior studies have demonstrated dendritic AVP release under hypovolemic conditions, direct confirmation, using sniffer-cell approaches or in vivo biosensors, will be needed to establish the mechanism with certainty. Second, while PEG is a well-established and widely used method to induce hypovolemia, the kinetics and magnitude of the hypovolemic state may differ from more physiological models such as controlled hemorrhage Thus, the extent to which these distinct protocols differentially engage AVP neurons or modulate central neurovascular responses remains to be tested. Third, the present study focused primarily on vascular diameter as the readout of neurovascular coupling. We did not assess whether PEG-induced hypovolemia also alters SON tissue oxygenation, blood flow velocity, or capillary perfusion, which would provide a richer physiological context and facilitate direct comparison with our previous salt studies and strengthen the mechanistic interpretation. Finally, the cellular basis for the differences between the osmotic and hypovolemic stimuli in the evoked NVC in HF rats remains unresolved. Thus, future studies testing pharmacological or genetic disruption of microglial purinergic signaling during hypovolemic stimulation would be valuable in determining whether this pathway is selectively recruited by osmotic stimuli.

In summary, although osmotic and hypovolemic challenges both engage AVP-dependent iNVC under physiological conditions, only osmotic stimulation recruits an alternative neurovascular program in heart failure. This stimulus-dependent reorganization of SON vascular signaling underscores the need to define how specific homeostatic inputs shape hypothalamic neurovascular dynamics and contribute to maladaptive neurohumoral drive in cardiovascular pathology.

## ACKNOWLEDGEMENTS

Grant support was provided by NIH R01HL162575 to JES and JAF; AHA 916907 to RKR. We also acknowledge support provided by the Center for Neuroinflammation and Cardiometabolic Diseases at GSU.

